# Climate change, biotic yield gaps and disease pressure in cereal crops

**DOI:** 10.1101/2022.08.12.503729

**Authors:** Muhammad Mohsin Raza, Daniel P. Bebber

## Abstract

Plant diseases are major causes of crop yield losses and exert a financial burden via expenditure on disease control. The magnitude of these burdens depends on biological, environmental and management factors, but this variation is poorly understood. Here we model the effects of weather on potential yield losses due to fungal plant pathogens (the biotic yield gap, Y_gb_) using experimental trials of fungicide-treated and untreated cereal crops in the UK, and project future Y_gb_ under climate change. We find that Y_gb_ varies between 10 and 20 % of fungicide-treated yields depending on crop, and increases under warmer winter and wetter spring conditions. Y_gb_ will increase for winter wheat and winter barley under climate change, while declining for spring crops because drier summers offset the effects of warmer winters. Potential disease impacts are comparable in magnitude to the effects of suboptimal weather and crop varieties.

## Introduction

Sustainable intensification of agriculture aims to increase food production without exacerbating environmental impacts, thereby avoiding the need to further expand agriculture into natural ecosystems to satisfy growing market demand ^1,2^. A key metric for intensification is the crop yield gap, which is the fractional difference between the potential yield in a region under irrigated or rainfed conditions and the average yield actually achieved by farmers ^1,3^. The yield gap depends on numerous factors including crop genotype, nutrient deficiency, water stress, solar radiation, growing season temperatures, management factors (e.g. reliance on manual labour) and the effects of weeds, pests and diseases ^1,3,4^. Yield gaps shrink with economic development, as wealthier countries are able to invest more in technology, training, fertilizer and crop protection, but tend toward 20% as further improvements become economically and ecologically undesirable ^5^.

While recent research has quantified the contribution of suboptimal crop genetics and management to yield gaps, biotic burdens like weeds, pests and diseases tend to be ignored ^3,5^. Expert opinion suggests that around one fifth to one third of crop production is lost to pests and diseases globally ^6^, but little is known about how these losses vary in time and space. Observed losses are potential losses reduced by expenditure on measures like weeding, disease-resistant seed, and agrochemical herbicides, pesticides and fungicides ^1,7,8^. Here, we focus on the impacts of fungal diseases. Disease risk varies with pathogen virulence, crop susceptibility and environmental factors like weather ^8,9^. Pest and disease life cycles are strongly determined by weather conditions, and many weather-driven models have been developed to predict occurrence or infection risk and thereby support decisions on when to apply control measures ^8^. Similarly, climate change, particularly warming, has driven historical increases in pest and disease incidence ^10^ and is likely to cause significant shifts in pest and disease risks in future ^11,12^. In contrast with disease risk, the effects of weather and climate change on yield losses to biotic agents are poorly understood.

Quantifying potential yield losses to biotic agents and why these vary is key to understanding an important component of crop yield gaps, and how to reduce them. The potential biotic yield gap (Y_gb_) can be defined as the fractional difference in yield between crops that have been protected against losses to biotic agents (Y_t_) and those that are unprotected (Y_c_) keeping crop variety and environment constant, i.e. 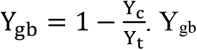 can be considered as a measure of disease pressure or disease burden, as it indicates the importance of disease to a particular cropping system. Potential losses can be estimated by controlled field experiments that compare protected (e.g. fungicide-treated) with control (untreated) yields. Such experiments are generally undertaken by agencies responsible for crop variety selection when determining pest or disease resistance levels, most often to fungal pathogens ^13^. Here, we analyse untreated (Y_c_) and fungicide-treated (Y_t_) yields from nearly two decades of grain cultivar trials in the UK to quantify Y_gb_ attributable to fungal pathogens (Supplementary Table 1), and to test the hypothesis that the fungal disease burden will increase with climate change. Further, we quantify the contribution of crop variety differences and interannual (climatic) variability to trial yields, and estimated the relative contributions of changing temperature and moisture to Y_gb_.

## Results

### Yields and the biotic yield gap

Yields varied among crops and between spring and winter varieties of wheat and barley (Fig. 1, Supplementary Fig. 1). Mean Y_t_ per site (averaged across all varieties) over the study period was 10.6 ± 1.7 (sample SD) t ha^-1^ for winter wheat, 7.2 ± 1.4 t ha^-1^ for spring wheat, 9.1 ± 1.4 t ha^-1^ for winter barley, 7.3 ± 1.2 t ha^-1^ for spring barley and 7.2 ± 1.5 t ha^-1^ for spring oats. Y_t_ tended to increase over time for winter and spring barley but not for the other crops. Mean Y_c_ per site was 8.3 ± 1.7 t ha^-1^ for winter wheat, 6.1 ± 1.2 t ha^-1^ for spring wheat, 7.4 ± 1.2 t ha^-1^ for winter barley, 6.6 ± 1.3 t ha^-1^ for spring barley and 6.3 ± 1.4 t ha^-1^ for spring oats. Y_c_ followed similar temporal trends to Y_t_, increasing for barley but not changing over the study period for the other crops. The mean difference between Y_t_ and Y_c_ for each individual variety trial was 2.3 ± 1.6 t ha^-1^ for winter wheat, 1.4 ± 1.2 t ha^-1^ for spring wheat, 1.7 ± 1.2 t ha^-1^ for winter barley, 0.8 ± 0.8 t ha^-1^ for spring barley and 1.0 ± 1.0 t ha^-1^ for spring oats. The mean biotic yield gap (Y_gb_) attributable to fungal pathogens per site was 0.21 ± 0.12 for winter wheat, 0.17 ± 0.13 for spring wheat, 0.18 ± 0.10 for winter barley, 0.11 ± 0.09 for spring barley and 0.13 ± 0.11 for spring oats. No trends were apparent in Y_gb_ over time for any crop.

**Figure 1.**
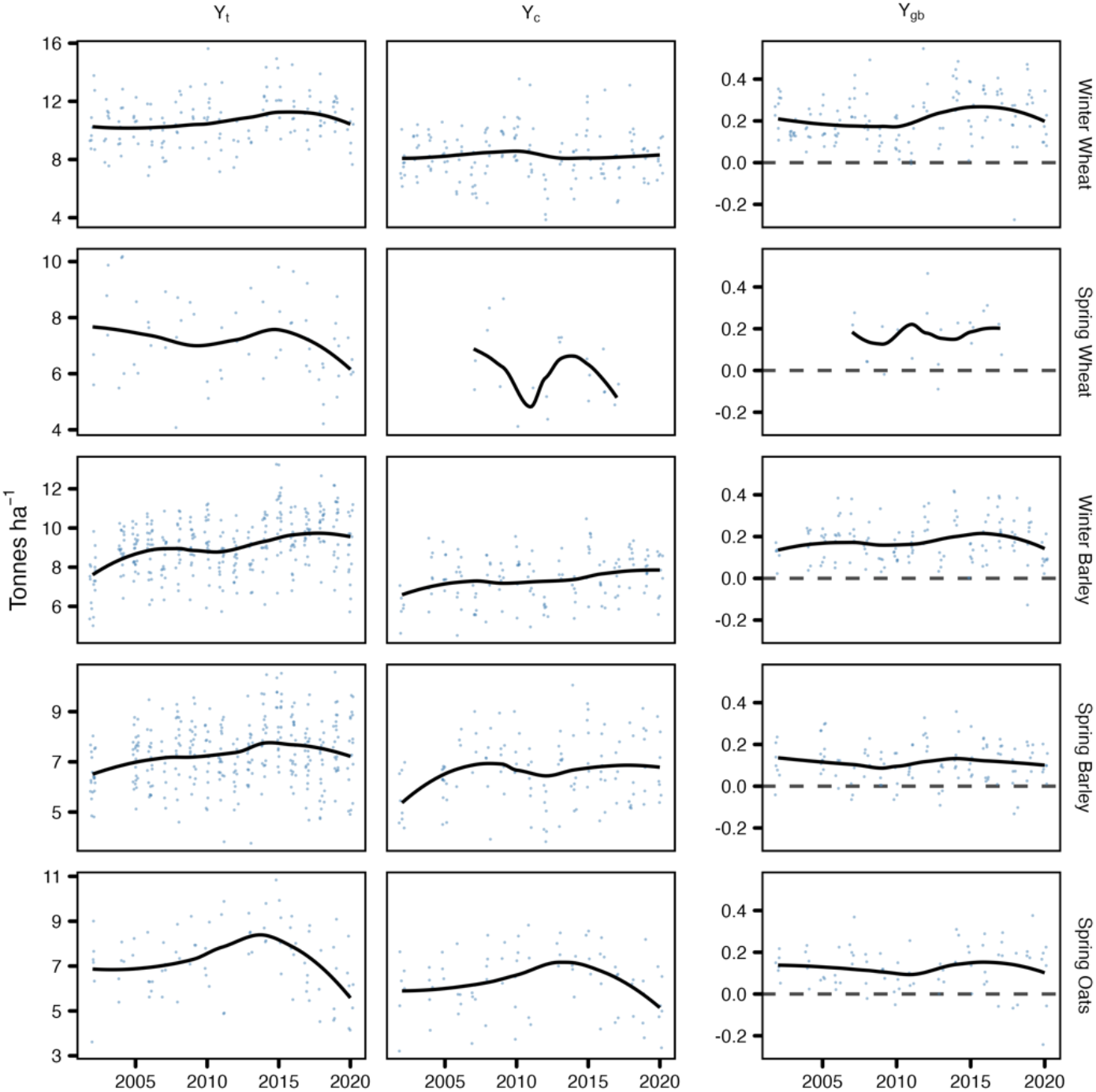
Distribution of total grain yield (Tonnes ha^-1^) and yield gap (Y_gb_). Distribution of grain yield from fungicide treated (Y_t_) and non-treated (Y_c_) experimental sites and resultant yield gaps (Y_gb_) in the studied crops from 2002 - 2020. The dashed horizontal line in the right panel indicates no yield gap. Data points above this line indicate yield gap (Y_t_ > Y_c_). However, points below this line indicate yield gain (Y_t_ < Y_c_).

### Maximum attainable yield and components of the yield gap

While Y_t_ estimates yield in the absence of fungal pathogens, the effects of genetic variation among varieties, growing season climate, and site-specific edaphic factors may reduce yield below what is potentially possible for a crop. We estimated the maximum attainable yield (Y_max_) for each crop from the top 5 % of all Y_t_ values across all trials. We detected no spatial trends in Y_t_ except for an increase with latitude in spring oats (Supplementary Table 2), and therefore estimated Y_max_ across all sites rather than as a function of location. Mean Y_max_ was 14.5 t ha ± 0.1 t ha^-1^ (bootstrap SD) for winter wheat, 10.1 ± 0.1 t ha^-1^ for spring wheat, 12.1 ± 0.05 t ha^-1^ for winter barley, 10.0 ± 0.03 t ha^-1^ for spring barley and 10.5 ± 0.1 t ha^-1^ for spring oats. We estimated the contribution of variety (genetic) differences to yield by the mean absolute error (MAE) of Y_t_ among varieties within sites and years (Y_gg_). Over the study period, Y_gg_ was 0.4 t ha^-1^ for winter wheat, 0.3 t ha^-1^ for spring wheat, 0.4 t ha^-1^ for winter barley, 0.3 t ha^-1^ for spring barley and 0.6 t ha^-1^ for spring oats. The MAE of Y_t_ within varieties and sites across years gave an estimate of the contribution of climatic variation to the yield gap (Y_gc_). Over the study period, Y_vc_ was 1.0 t ha^-1^ for winter wheat, 0.7 t ha^-1^ for spring wheat, 0.7 t ha^-1^ for winter barley, 0.6 t ha^-1^ for spring barley and 0.7 t ha^-1^ for spring oats.

Taking winter wheat as an example, we decomposed the gap between Y_max_ and Y_min_ (the mean of the lowest 5 % Y_c_ values) into biotic (Y_gb_), genetic (Y_gg_) and climatic (Y_gc_) components (Fig. 2). In this case, Ygg and Ygc were the empirical 95 % confidence intervals of Y_t_ deviations rather than MAE, indicating the difference between best and worst varieties within trials, and best and worst years within varieties. This indicated that mean losses to disease were of similar magnitude to varietal effects, but smaller than the effects of interannual climatic variation. Modelled potential yields and achieved yields for rainfed wheat^14^ lie within the range of Y_max_ and Y_min_.

**Figure 2.**
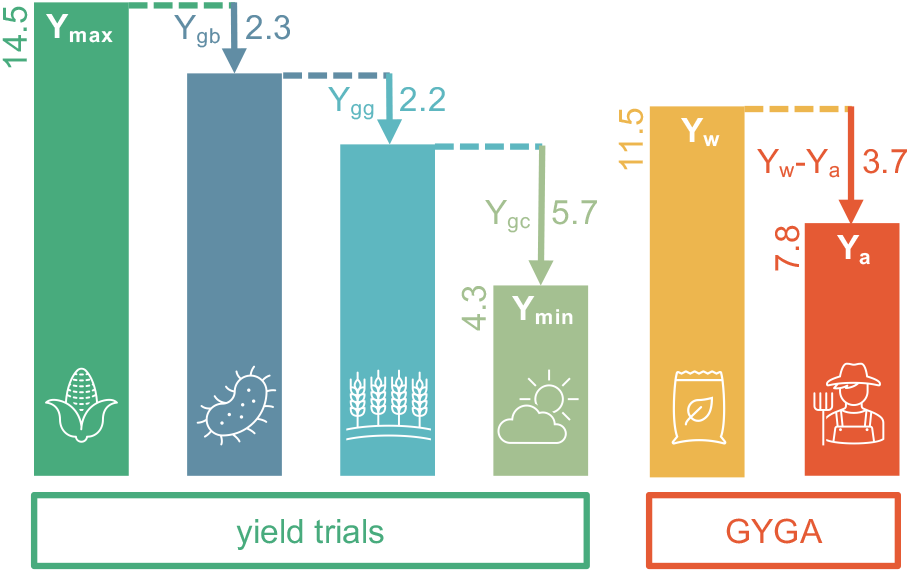
Yield gap components in winter wheat. Y_max_ and Y_min_ are top and bottom 5 % of Y_t_ and Y_c_, respectively, across all trials. Y_max_ (14.5 t ha^-1^) indicates the highest yield achievable under optimal weather conditions in the best sites with the best varieties and no loss to disease. Y_min_ (4.3 t ha^-1^) indicates the yield in the worst sites with the lowest yielding varieties in the worst years with large losses to disease. Y_gb_ (2.3 t ha^-1^) is the mean loss to disease. Y_gg_ (2.2 t ha^-1^) indicates the difference between the highest and lowest Y_t_ of varieties within trials. Y_gc_ (5.7 t ha^-1^) indicates the difference between the best and worst Y_t_ of a variety within a site. Yield trial results are compared to modelled potential rainfed wheat yield for the UK (Y_w_) and achieved yield (Y_a_) from the Global Yield Gap Analysis^14^. Bar are not to scale.

### Weather and the biotic yield gap

We correlated Y_gb_ with site-specific monthly temperature, relative humidity (RH) and precipitation over the growing season to determine the most important weather variables driving fungal disease pressure (Fig. 3). Winter temperatures and summer RH were most strongly positively correlated with Y_gb_ in winter wheat and in barley, while spring and summer precipitation were most important in spring wheat. Early spring temperature and early summer RH were most strongly correlated with Y_gb_ in spring oats. We selected the single months with the strongest temperature and RH (or precipitation) correlations for each crop for predictive modelling. The correlations for the best predictor months varied between 0.22 and 0.57 (Supplementary Table 3). Inclusion of additional months in the models was unnecessary because weather is temporally autocorrelated (a warmer February tends to follow a warmer January etc). Model selection determined that, over the range of monthly temperature and humidity values in the data, the relationships with Y_gb_ were best explained by additive linear terms, except for spring oats for which there was an interaction between March temperature and May humidity (Supplementary Fig. 2, Supplementary Table 4). Fitted values for the models were strongly correlated (r > 0.41) with observations (Supplementary Table 4).

**Figure 2.**
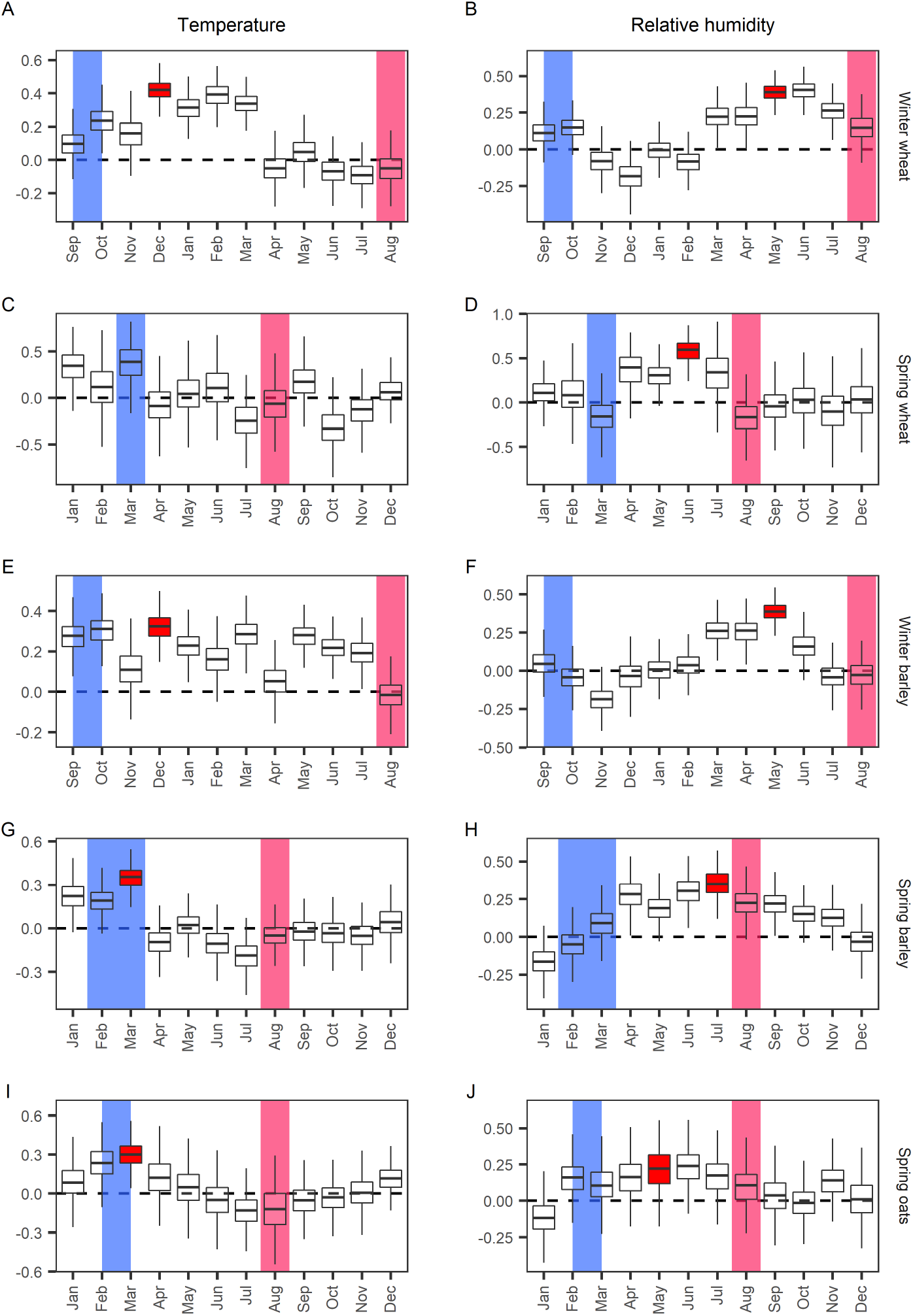
Bootstrapped estimates of the correlation between yield gap (Y_gb_) and monthly weather variables. Boxplots show the distribution (minimum, maximum, median and interquartile range) of correlation coefficient (*r*) estimates of the association between Y_gb_ and monthly temperature and relative humidity except for D, representing the correlation estimates between Y_gb_ and rainfall. Dashed horizontal line indicates no correlation. Blue and pink shaded areas indicate planting and harvesting times of the studied AHDB crops, respectively. The varying pattern of boxplots indicates how the correlation estimates vary for weather variables in each month of the growing season. Boxplots filled with red are the months we used climate data for model fitting. We did not find any significant association of Y_gb_ with temperature in spring wheat (C).

### Climate change and the biotic yield gap

We estimated Y_gb_ across crop production areas in the UK with our models, under recent historical (2002 – 2020) and projected future climates (2021 – 2040 and 2061 – 2080). Wheat production is currently concentrated in central and eastern England, barley in central southern England and eastern England and Scotland, and oat production occurs at low densities across the country (Supplementary Fig. 3). We employed the Met Office UKCP18 RCP8.5 projections at 5 km resolution for both the historical and future climates. We used current crop distributions for all estimates and did not try to project potential future crop distributions. Mean Y_gb_ weighted by crop area was around one fifth for winter wheat, spring wheat and winter barley, and one tenth for spring barley and spring oats over the recent historical period (Fig. 4, Supplementary Table 5). For all crops Y_gb_ increased towards the South and West (Fig. 4). On average, Y_gb_ increased slightly in the two future periods for winter wheat and winter barley, but declined slightly for the spring crops (Supplementary Table 5). Our results were robust to model perturbations in future climate projections, with mean standard deviations below 0.015 % across the production areas of each crop (Supplementary Figs. 4 – 8, Supplementary Table 6).

**Figure 4.**
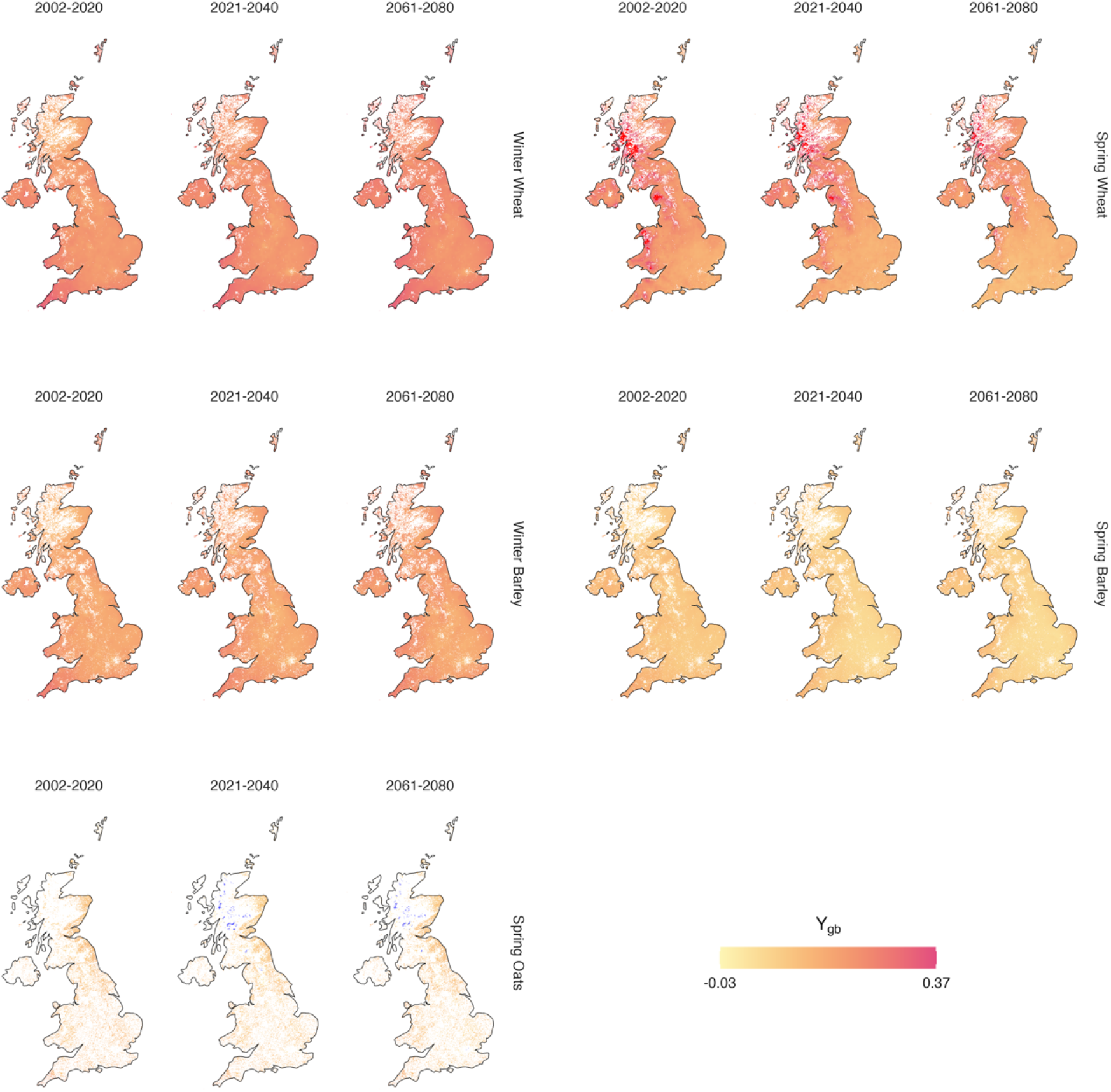
Current and future predicted average yield gap (Y_gb_). Predictions were made on current (2002 - 2020) and future (2021 – 2040 and 2061 - 2080) climate pixels of 1km x 1km grid resolution. Predicted Y_gb_ were then summarized for each time slice. Grey indicates either no or low Y_gb_, while dark blue represents a higher Y_gb_. White grid cells contained no hosts and were excluded from the analysis. Values outside 1.5 times the interquartile range (IQR) above upper quartile and below lower quartile are shown in red and deep blue respectively.

There were marked spatial patterns in the projected future changes in Y_gb_ (Fig. 5). For winter wheat and winter barley, Y_gb_ tended to increase in Wales, East Anglia, northern England and Scotland. For spring wheat the greatest declines in Y_gb_ were projected in the South West while small increases occurred in Scotland. For spring barley, the change in Y_gb_ was projected to be negative over most of England and Wales, and positive in northern Scotland. Spring oats are less commonly planted across the UK, but there was some indication of an increase in Y_gb_ in Wales with declines elsewhere. Overall, our results suggest that on average the changes in Y_gb_ will be relatively minor, but that some regions will experience large increases or decreases in fungal disease pressure depending on the crop. In particular, spring crops will see overall decreases in Y_gb_, while winter crops will see increases. This difference between winter and spring crops is attributable to projected changes in temperature and moisture in winter and summer (Supplementary Figs. 9 – 10). Winter temperatures (December to February) increase less than summer temperatures (June to August), with the largest increases expected in the south. Winters are expected to get wetter, particularly in the north, while summers are expected to become drier in the south and wetter only in northern Scotland. Most winter wheat and barley production occurs in regions that will warm substantially in December and become drier in May (Fig. 6). These trends have opposing effects on Y_gb_, meaning that much of the production area is expected to experience relatively small changes in Y_gb_. In contrast, most of the production area for spring barley occurs in areas expected to experience only moderate March warming, but substantial drying in July. This results in declines in Y_gb_ for the majority of the production area. Most of the production area for spring oats is expected to experience only moderate changes in temperature and moisture, with relatively minor associated changes in Y_gb_.

**Figure 5.**
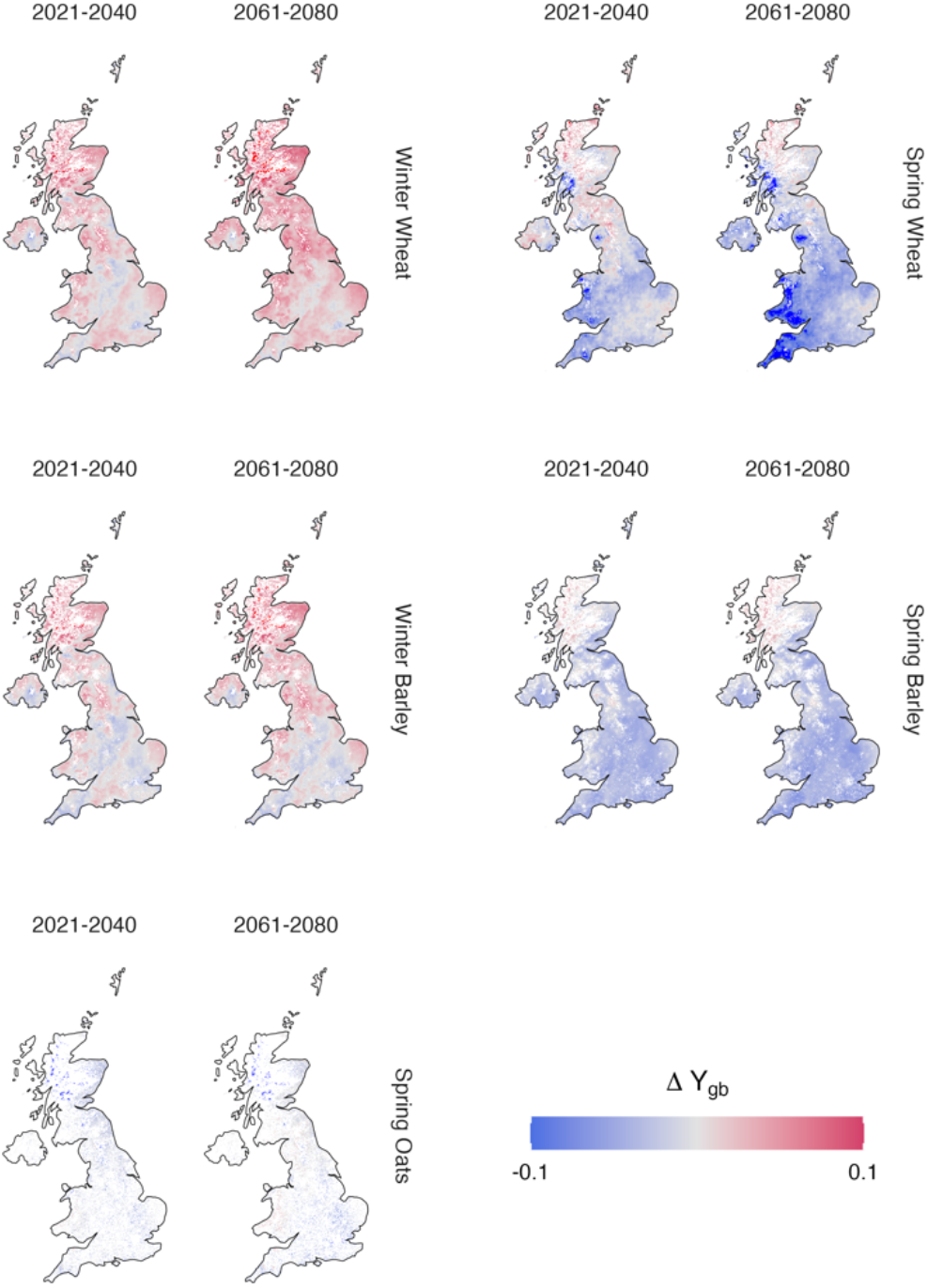
Average change in future yield gap (ΔY_gb_). Current (2002 - 2020) average predicted Y_gb_ levels were subtracted from future (2021 – 2040 and 2061 - 2080) predicted Y_gb_ levels for each pixel at 1km x 1km grid resolution. Red indicates a high Y_gb_, while blue indicates high yield gain compared to current Y_gb_ levels. Grey indicates no change. White grid cells contained no hosts and were excluded from the analysis. Values outside 1.5 times the interquartile range (IQR) above upper quartile and below lower quartile are shown in red and deep blue respectively.

**Figure 6.**
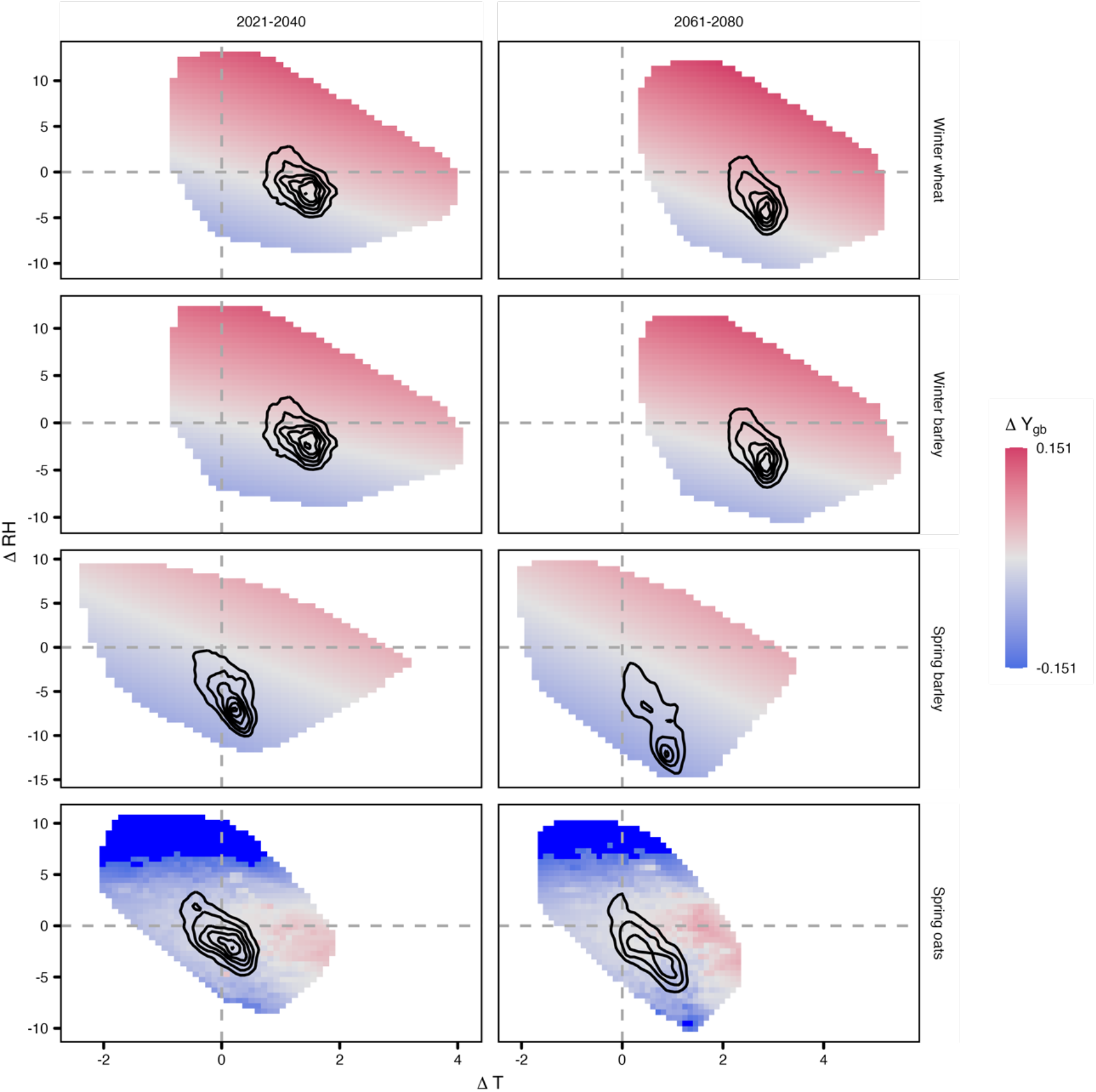
Association between change in average future temperature (ΔT) and relative humidity (ΔRH) and the interpolated surface of mean yield gap differences (ΔY_gb_) in future time slices. Contour lines represent the aggregated count of data points of association between ΔT and ΔRH. Values outside 1.5 times the interquartile range (IQR) below lower quartile are shown in deep blue.

## Discussion

Our results show that fungal disease pressure on grain crops in the UK, as measured by Y_gb_, amounts to between one tenth and one fifth of yield in variety trials. Y_gb_ tends to increase with winter temperatures and summer moisture, and Y_gb_ is greater in wheat and in winter barley than in spring barley and spring oats. Projections of Y_gb_ under future climates using these models suggests that change in disease pressure will be moderate on average, but spatially variable and dependent upon the crop growing season. Winter varieties are more likely to see increases in disease pressure due to warming winters, which are only partially offset by drying summers. Spring varieties of wheat and barley are likely to see declines in disease pressure due to summer drying. These changes could influence the relative importance of spring and winter varieties in the UK in future. While our models cannot be reliably extrapolated outside the UK, the strong predictive power of our relatively simple models suggests that our approach could be applied to other regions where suitable agricultural trial data are available.

We did not attempt to relate Y_gb_ to incidences of specific fungal diseases. AHDB provides disease incidence scores for a number of pests and pathogens for each crop, but available records are highly incomplete making statistical estimation of impacts difficult. Septoria Tritici Blotch (STB, caused by *Zymoseptoria tritici*) has been the most important disease of winter wheat in the UK for several decades ^15^. Other significant fungal diseases of winter wheat include brown rust (caused by *Puccinia triticina*), yellow rust (*Puccinia striiformis*), the soilborne disease take-all (*Gaeumannomyces tritici*), glume blotch (*Phaeosphaeria nodorum*), powdery mildew (*Blumeria graminis*), tan spot (*Pyrenophora tritici-repentis*), eyespot (*Oculimacula* spp.), sharp eyespot (*Rhizoctonia cerealis*), and Fusarium ear blight (*Fusarium* spp.) ^15^. Farm surveys between 1999 and 2019 suggest that incidences of most diseases are rather variable over time, with glume blotch, powdery mildew, eyespot and sharp eyespot declining somewhat and Fusarium ear blight emerging ^15^. Temporal and spatial dynamics of fungal diseases of spring wheat, barley and oats are less well characterized than those of winter wheat.

Fungal plant pathogens show a range of climatic tolerances ^16^, therefore the suite of diseases affecting different crops may well change in future with warming and other global change drivers ^12^. For example, improvements in air quality in recent decades may have allowed STB to overtake glume blotch as the most important winter wheat disease in the UK ^17^, although changes in fungicide application regimes are also implicated ^15^. A combination of climate change, landscape management and crop breeding may allow a previously important disease, stem rust (caused by *Puccinia graminis* f.sp. *tritici*), to return ^18^. Our projections of future Y_gb_ assumed that fungal disease responses to weather would remain constant, though this may not be tenable if the pathogen assemblage changes. However, general trends in fungal pathogen responses to climate change have been reported. For example, soilborne fungal pathogens tend to increase in relative abundance in response to warming ^19^. Replication of our methods in other regions, thereby extending the climate envelope for model parameterization, could help to determine the generality of the patterns we have detected.

We assumed that fungicide applications in trials completely prevented yield losses. In the UK, fungicides are applied to nearly all crop areas with between three and four sprays applied to winter wheat during the growing season ^15^. The most consistently important fungicide classes have been demethylation inhibitors. Use of strobilurins has declined due to resistance evolution, while succinate dehydrogenase inhibitors and chlorothalonil use has increased ^15^. Details of experimental fungicide applications are not reported by AHDB, but we assumed that the manufacturer-recommended dosage and timings were implemented. Farmers tend to apply less than the recommended dosage, though this fraction increased from around 0.4 to around 0.8 between 1999 and 2019 ^15^. Fungicides are required because genetic resistance to fungal pathogens provides insufficient protection. Resistance to STB and Wheat mildew (*Blumeria graminis* f. sp. *tritici*) is polygenic and partial, but tends to be durable over time, while resistance to rusts and Barley mildew (*Blumeria graminis* f. sp. *hordei*) is monogenic and persists for a few years before being overcome by evolution of virulence in the pathogen ^20^. Variation in resistance will be a major determinant of the variability in Y_gb_ among tested varieties.

We statistically modelled Y_gb_ in relation to weather while the majority of studies have focussed on processes like infection rate or some measure of disease risk ^8,12,21–23^. Process-based, or mechanistic, models of infection risk tend to be driven by hourly meteorological data ^22^, though some large-scale studies have employed monthly averages ^12^. Temperature responses are usually humped, with the maximum infection rate occurring at optimum temperature. In contrast, we detected a linear response to temperature. This may indicate that UK crop production occurs at temperatures below the optima for important fungal pathogens. The effect of moisture is commonly modelled as an increasing function of humidity, or a binary process whereby infection can only take place during periods in which leaf surfaces are wet ^22^. While these models of disease risk can be used in disease control decision-making, or to estimate risks under future climates, they do not directly estimate potential yield losses. In the UK, potential yields of rainfed crops (Y_w_) estimated from crop models ^5,14^ vary between 11.2 and 12.9 (mean 11.5) t ha^-1^ for wheat and 8.5 and 9.8 (mean 8.9) t ha^-1^ for barley, depending on climate zone ^5^. Achieved yields (Y_a_) vary between 7.3 and 8.1 t ha^-1^ (mean 7.8 t ha^-1^) for wheat and 5.6 and 6.3 t ha^-1^ (mean 6.0 t ha^-1^) for barley. Oats are not modelled, and winter and spring varieties are not differentiated ^5^. The modelled yield gap between Y_w_ and Y_a_ is therefore 3.7 t ha^-1^ for wheat and 2.9 t ha^-1^ for barley. While we cannot estimate the contribution of different causes (weather, variety selection, pests and diseases) precisely, our results demonstrate that potential losses from pathogens are a similar magnitude to other yield gap drivers (Fig. 2), and that climate change will differentially affect varieties and could therefore influence cropping patterns.

## Methods

### Crop and Yield data

We analysed yield data for crop variety trials conducted by the Agriculture and Horticulture Development Board (AHDB) from 2002 to 2020. AHDB hosts archives of recommended lists of cereals and oilseed that provide independent information on yield and quality performance, agronomic features, disease pressure and market options to assist with variety selection ^24^. This list is updated each year and provides information based on the analysis of hundreds of UK trials conducted since 2002. No information is provided for fungicide usage in trials. We also obtained the approximate spatial coordinates for the trial locations for data analysis and mapping by matching names of trial locations to locations using GeoNames ^25^ and Google map search (Supplementary Fig. 11). Data were cleaned to remove sites and varieties with missing data for yields. Yield from fungicide-treated (Y_t_) and untreated (Y_c_) trials was used to calculate the fungal disease-related yield gap (Y_gb_) as:

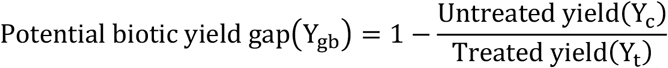

As each site had data for different varieties and cultivars, Y_gb_ and disease pressure information, mean values per site per year were used for subsequent analyses.

Fungicide-treated (Y_t_) and untreated (Y_c_) yields were available for winter wheat (289 varieties), spring wheat (47), winter barley (147), spring barley (154) and spring oats (45). Site locations varied geographically between Limavady, Northern Ireland (6.98 °W, 55.07 °N) in the west and Morley, Norfolk (1.03 °E, 52.56 °N) in the east, and West Charleton, Devon (3.76 °W, 50.27° N) in the south and Kinghorn, Fife (3.10° E, 56.18° N) in the north (Supplementary Fig. 1). The number and locations of trial sites varied over time, and the number and composition of varieties in the trials varied among sites and over time. The total number of crop varieties tested per year remained roughly stable, with a mean of 42 winter wheat, 9 spring wheat, 22 winter barley, 22 spring barley, and 10 spring oat varieties tested per year.

### Crop map

Crop distribution data for the UK were obtained from the EUCROPMAP 2018 ^26,27^. This map is produced using Sentinel S1A and S1B Synthetic Aperture Radar observations for 2018 and random forest-based classification algorithms. The map provides detailed spatial information on 19 crop types in the EU for 2018 at 10-m resolution, with high accuracy. Pixels for each AHDB crop were extracted and aggregated to a 1 km x 1 km grid generated using the *Fishnet* tool in ArcMap (ESRI ArcGIS Desktop, release 10.8). This grid was generated on the same 1 km grid as the HadUK-Grid Gridded Climate Observations. The count of crop pixels in each grid cell was then used to calculate the total area (hectares) of crop under cultivation and, subsequently, the fractional area under crop cultivation (A_F_) in each 1 km grid cell for further analysis (Supplementary Fig. 3).

### Climate data

Monthly weather data (2001-2020), including mean air temperature (°C), relative humidity (%), total rainfall (mm) and sunshine hours (h), were obtained from the HadUK-Grid Gridded Climate Observations v1.0.3.0 on a 1km and 5km grid over the UK ^28^. Weather data from 1km grid were extracted for AHDB trial sites (point locations, years 2001 - 2020) for each crop from the rasters using the *extract* function of the *raster* package in R v. 4.2.1 ^29^. Climate model projections for monthly mean air temperature (°C), relative humidity (%), total rainfall (mm) and sunshine hours (h) were obtained from UKCP Local Projections on a 5km grid over the UK for 2021-2040 and 2061-2080 ^30^. These projections are produced by the Met Office Hadley Centre as part of the UK Climate Projection 2018 (UKCP18) project and cover three time-periods (1981-2000, 2021-2040 and 2061-2080) for a high emissions scenario, RCP8.5. Projections for other emission scenarios are not available. For Y_gb_ predictions, monthly weather data (2001-2020) from HadUK-Grid Climate Observations and climate model projections data (2021-2040 and 2061-2080) from UKCP Local Projections on a 5 km grid were extracted for areas under cultivation for each crop, using the *mask* function of *raster* package in R ^29^.

### Climate-yield gap relationship estimation

The relationship between Y_gb_ and climatic variables for each month was explored using simple correlation and regression analysis. Estimates from correlation and regression analyses were bootstrapped (1000 iterations) using the *bootstraps* function in the *rsample* package of R ^31^, where each iteration was fitted on a resampled dataset with replacement. We used temporal block bootstrapping to randomly resample data from a single year with replacement instead of sampling random monthly observations, to maintain within-year temporal correlations ^32^. The bootstrapped estimates of correlation coefficient and beta slope estimate were visually inspected for consistency and strength of association between monthly weather variables and observed Y_gb_.

Our objective was to statistically fit the observed Y_gb_ to monthly weather variables. The resulting relationship would be used for further analyses. We used generalized-least-squares (GLS) using the *gls* function from the *nlme* package in R ^33^ to fit the model between Y_gb_ and the significant weather variables identified through bootstrapping estimates. The GLS regression allowed for a first-order autoregressive correlation structure in the residuals to account for the correlation over time and among experimental sites in climate data. The parameters of the GLS regression represented the climate-driven trends in yield gaps. We compared the models with and without temporal correlation structure, using *F-test* in the *anova* function, to determine whether inclusion of temporal autocorrelation was required. Finally, current (2002-2020) and future yield gap levels (2021-2040, 2061-2080) were predicted and the estimates of the standard error of prediction were calculated for each masked (crop pixels only) climate dataset pixel (5 km) using the *predictSE*.*gls* function from the *AICcmodavg* package in R ^34^. The modelled yield gaps were averaged for each time period and climate dataset pixel. In addition, we made predictions on all 12-member perturbed physics ensembles (UKCP local 5 km) for projections to get uncertainty in Y_gb_ predictions due to climate model parameters/physics perturbations ^10^.

### Future climate change impacts on yield gaps (forecast) and climate risk classification

The impact of future climate change on Y_gb_ was quantified as the change in predicted future yield gaps (2021-2040, 2061-2080) relative to current yield gaps (2002-2020). The mean yield gap differences ΔY_gb_ were calculated for each pixel (5 km), weighted by A_F._ We also compared the relationship between ΔY_gb_ and the change in average future temperature (ΔT) and relative humidity (ΔRH) levels in each future time slice to identify the strong drivers of Y_gb_ in each crop.

## Data availability

Crop yield trial data were obtained from the Agriculture and Horticulture Development Board (AHDB); part of this dataset is from the AHDB Recommended Lists. The AHDB Recommended Lists are managed by a project consortium of AHDB, BSPB, MAGB and UKFM. Data are available from https://ahdb.org.uk/rl. All datasets used in the study are freely and openly available from sources described in the Methods.

## Acknowledgements

This work was supported by Wave 1 of The UKRI Strategic Priorities Fund under the EPSRC Grant EP/W006022/1, particularly the “Environment and Sustainability” theme within that grant & The Alan Turing Institute.

## Authors contributions

DB designed the study. MR analysed the data and prepared the figures. Both authors wrote the manuscript.

