## Supplementary Figures for "Climate change, biotic yield gaps and disease pressure in cereal crops"

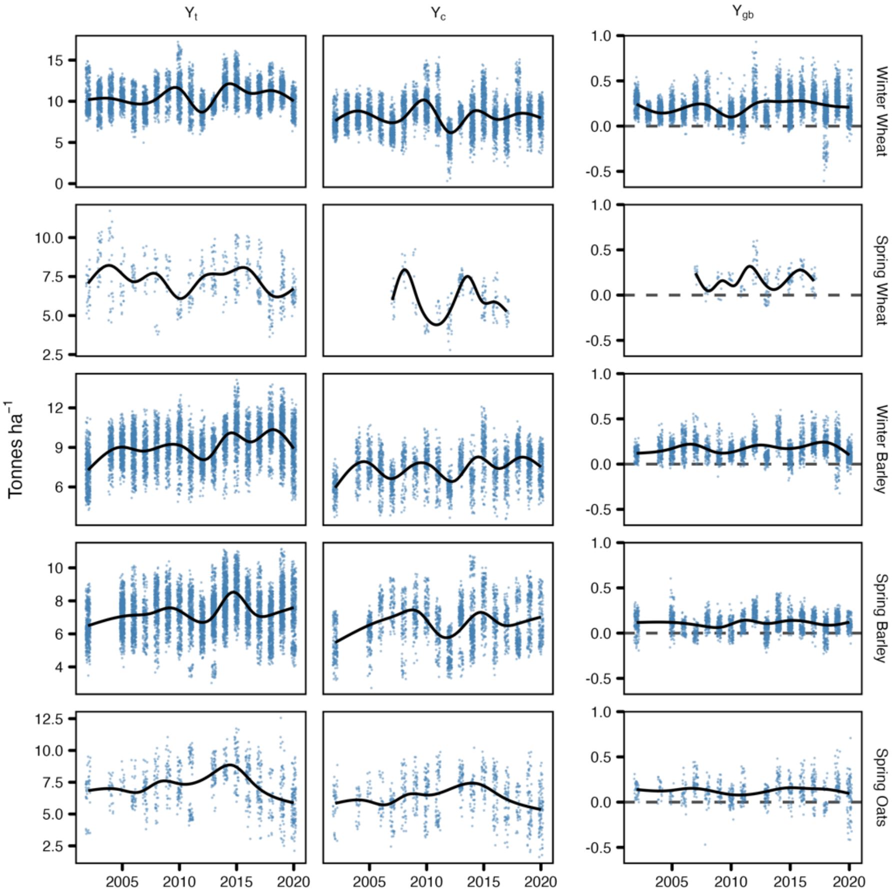


**Supplementary Fig. 1 |** Distribution of total grain yield (tonnes ha$^{-1}$) and yield gap ($Y_{gb}$) in AHDB studied crops. Points show results for individual varieties.

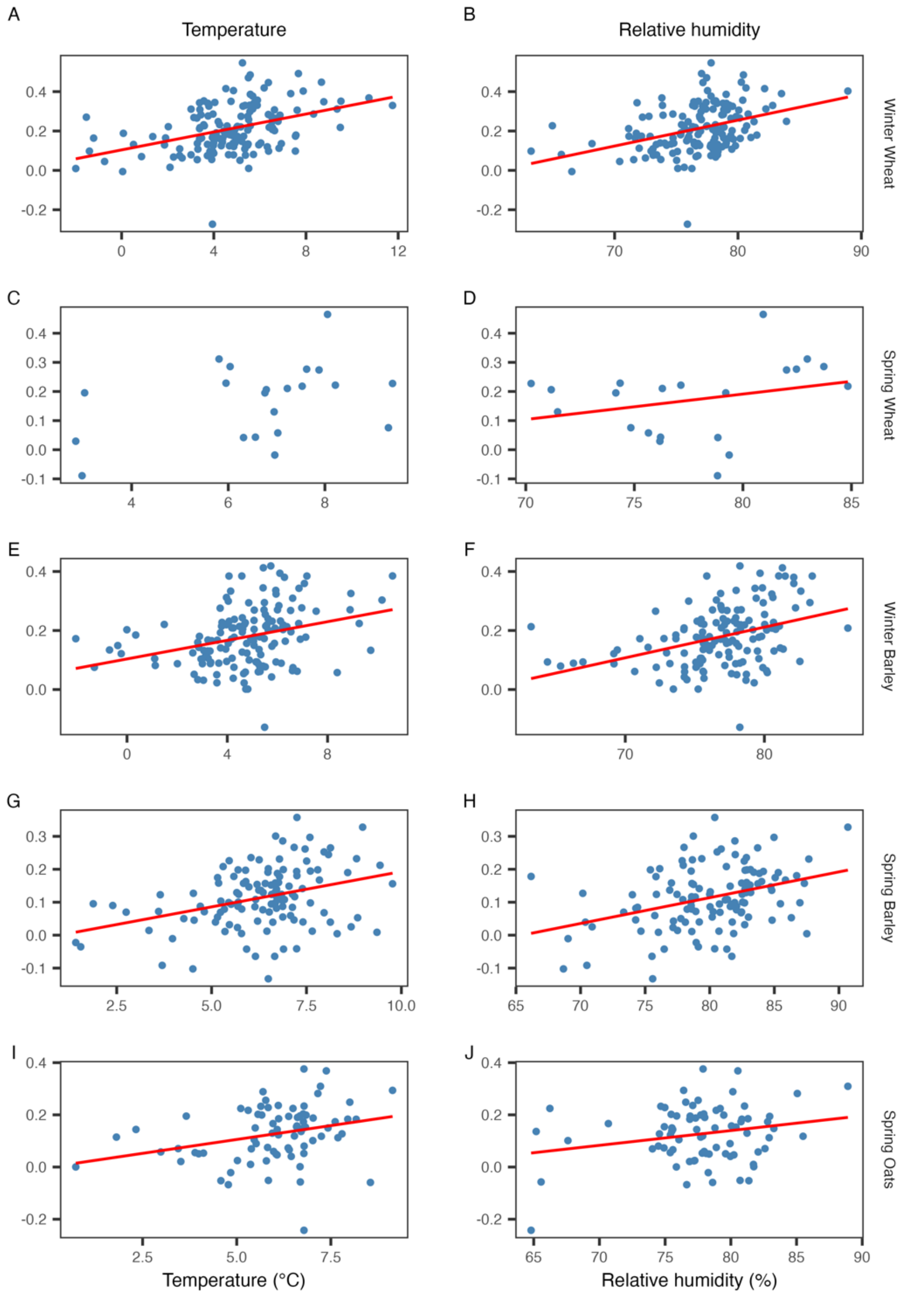


**Supplementary Fig. 2|** Scatter plots of the association of biotic yield gap ($Y_{gb}$) with monthly temperature and relative humidity except for D, representing the correlation estimates between $Y_{gb}$ and rainfall. We did not find any significant association of $Y_{gb}$ with temperature in spring wheat (C).

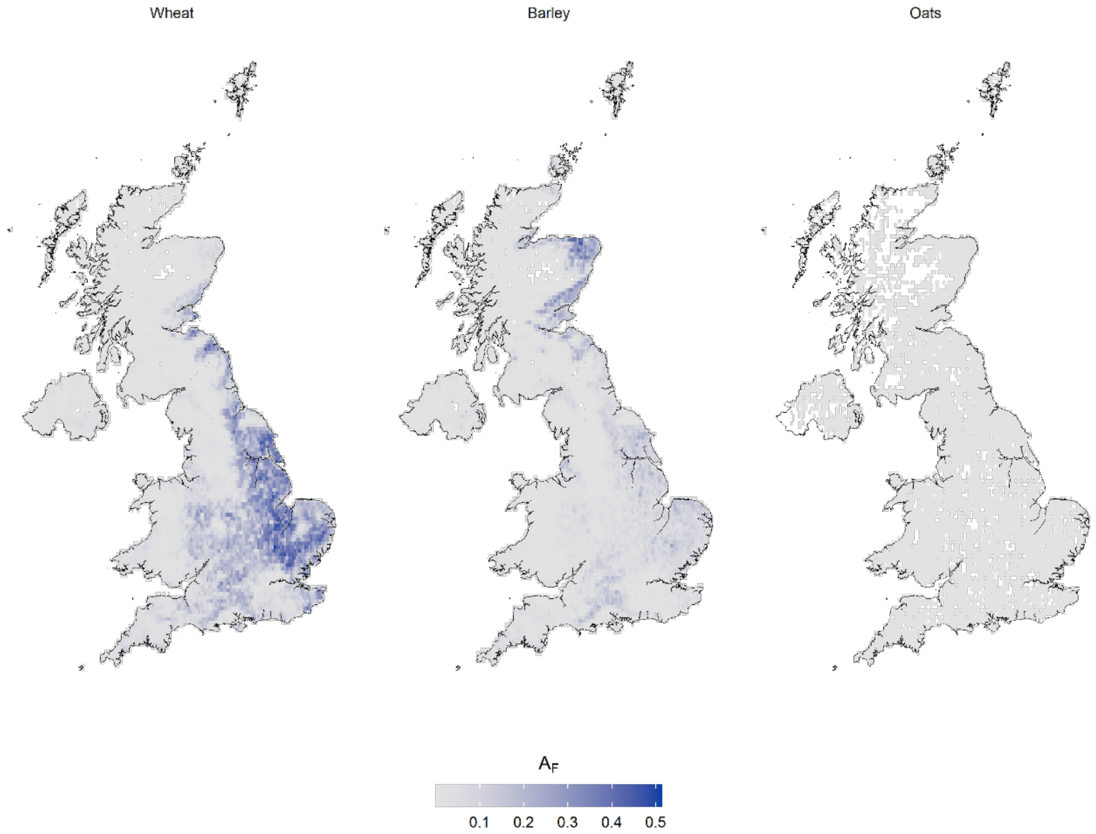


**Supplementary Fig. 3 |** Fraction area ($A_F$) under Wheat, Barley and Oats cultivation in the United Kingdom. Pixels (10-meter resolution) from crop type map were aggregated in 1km x 1km grids to calculate the area under cultivation as hectares. Fraction area was then calculated as the ratio of area under cultivation to the 1km x 1km grid area. Here, $A_F$ in 1km x 1km grids is further aggregated to 5km x 5km grids for better visualization. White grid cells contained no hosts and were excluded from the analysis.

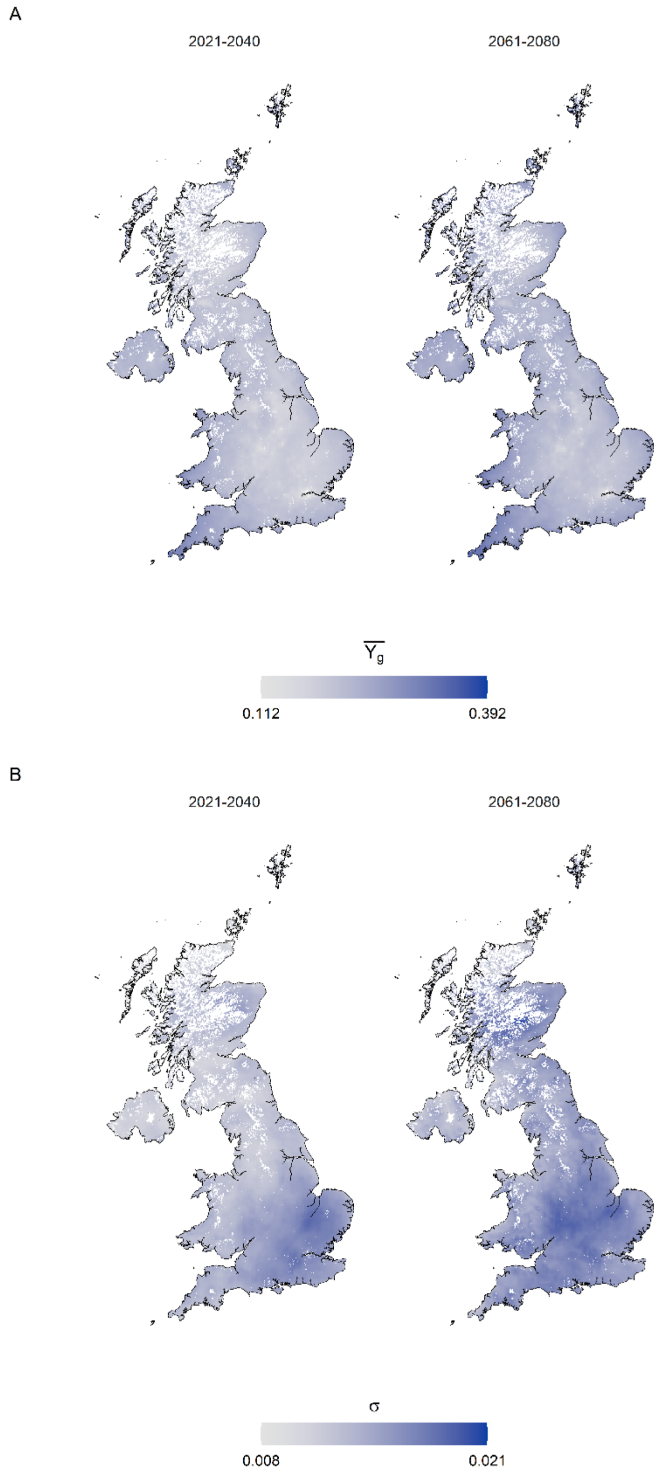


**Supplementary Fig. 4 |** Predicted future average yield gaps $Y_{gb}$ (**A**) in winter wheat and their standard deviations σ (**B**) using 12 physics perturbations of climate model projections from UKCP Local Projections on a 5km grid. White grid cells contained no hosts and were excluded from the analysis.

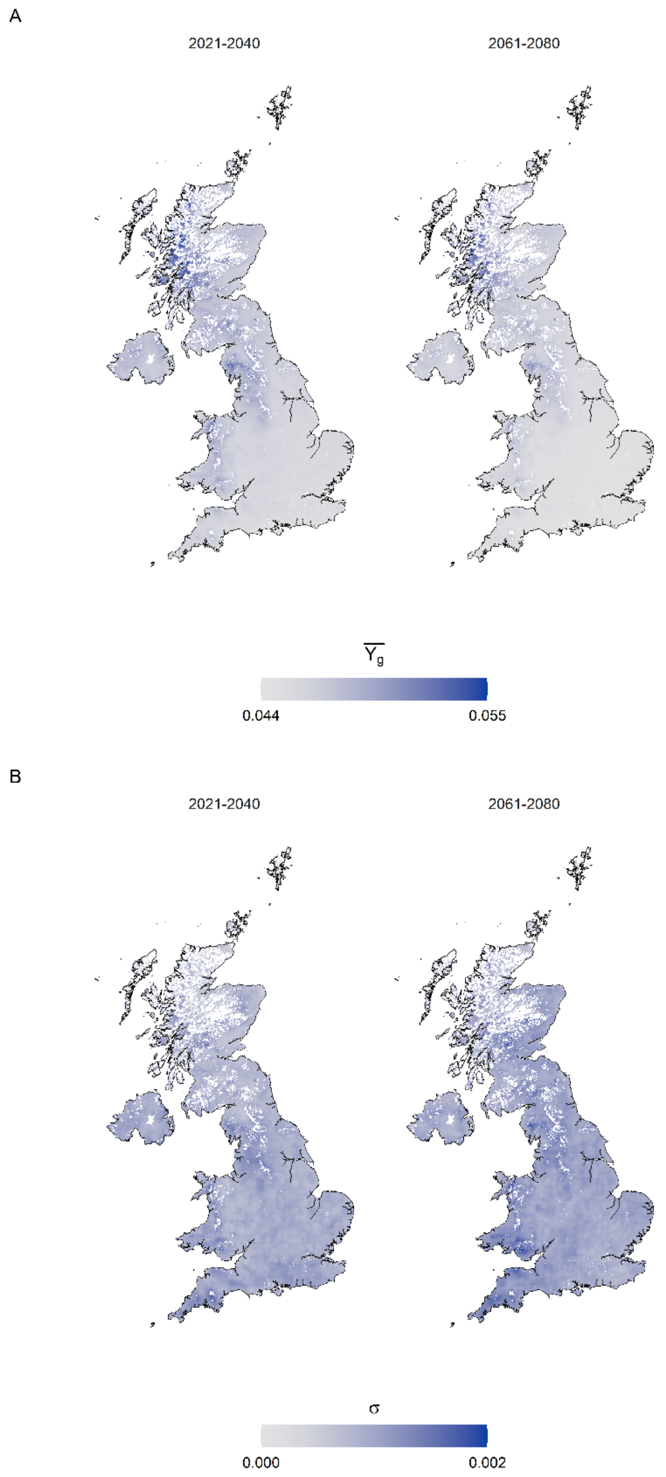


**Supplementary Fig. 5 |** Predicted future average yield gaps $Y_{gb}$ (**A**), in spring wheat and their standard deviations σ (**B**) using 12 physics perturbations of climate model projections from UKCP Local Projections on a 5km grid. White grid cells contained no hosts and were excluded from the analysis.

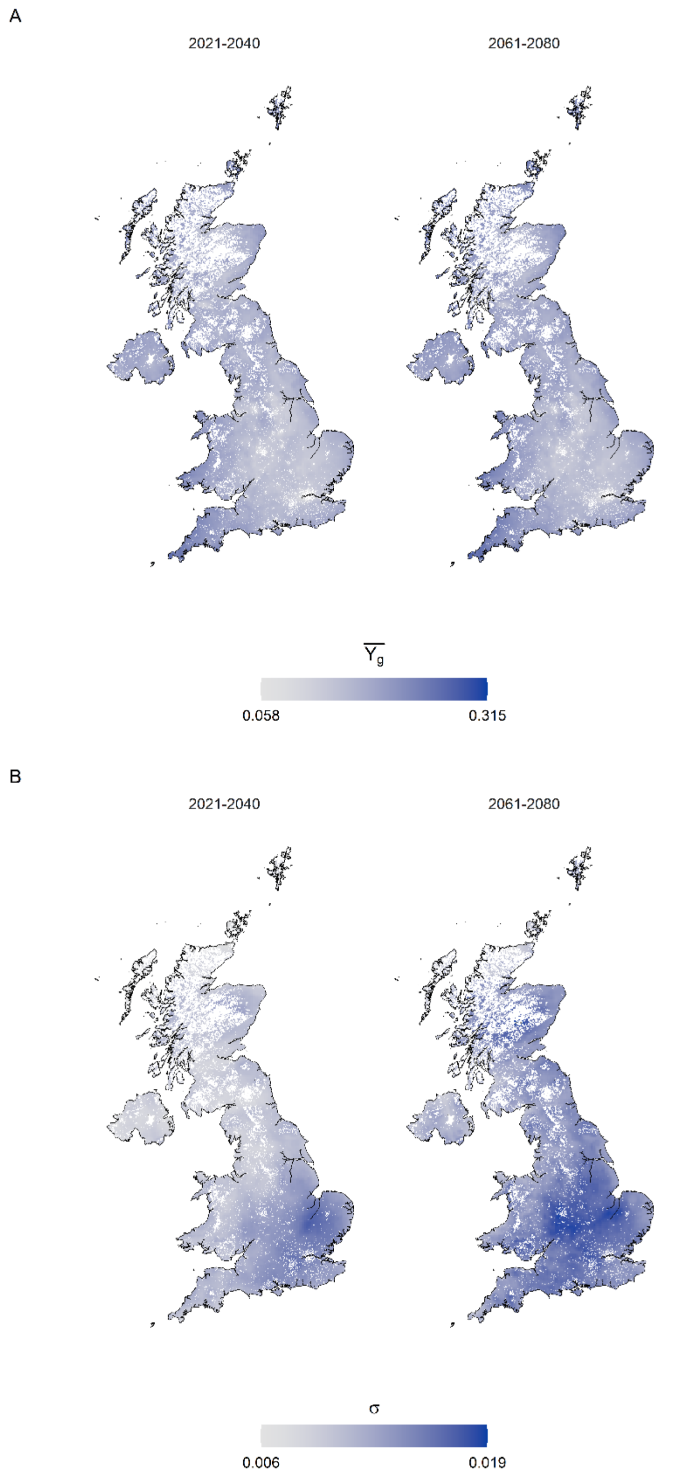


**Supplementary Fig. 6 |** Predicted future average yield gaps $Y_{gb}$ (**A**) in winter barley and their standard deviations σ (**B**) using 12 physics perturbations of climate model projections from UKCP Local Projections on a 5km grid. White grid cells contained no hosts and were excluded from the analysis.

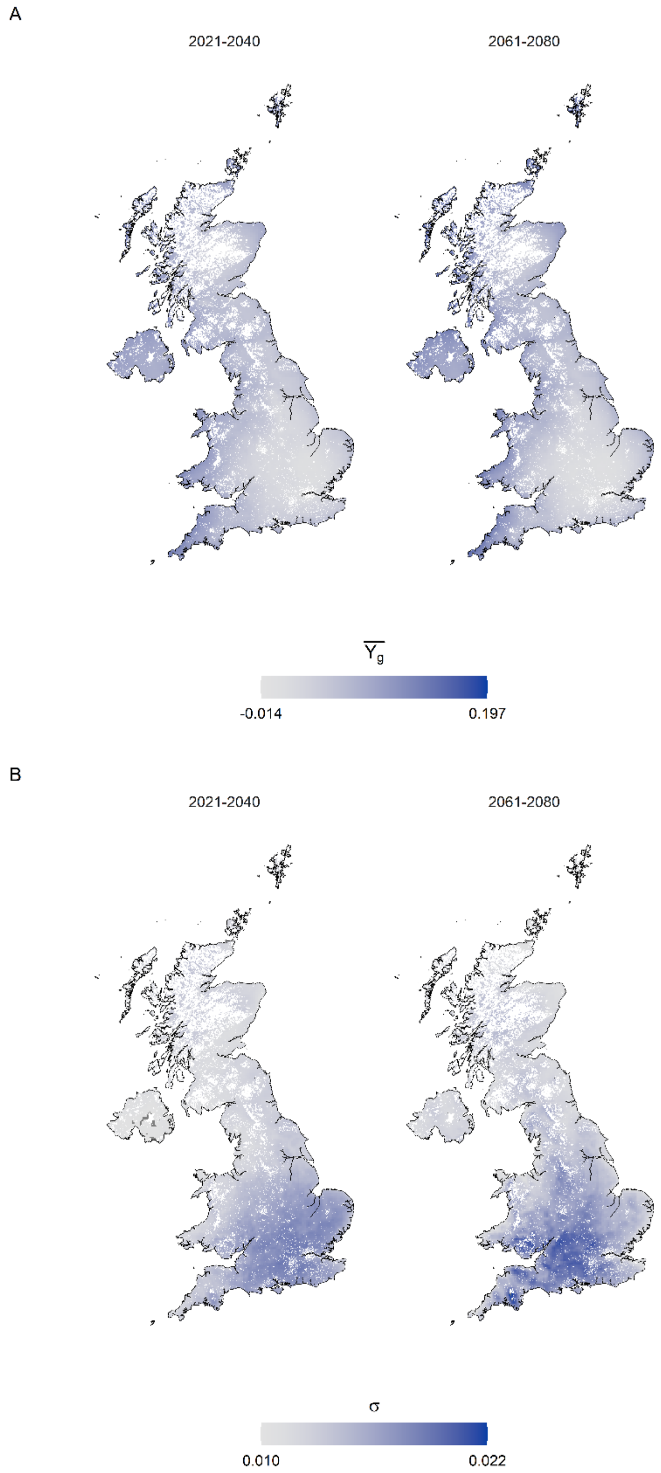


**Supplementary Fig. 7 |** Predicted future average yield gaps $Y_{gb}$ (**A**) in spring barley and their standard deviations σ (**B**) using 12 physics perturbations of climate model projections from UKCP Local Projections on a 5km grid. White grid cells contained no hosts and were excluded from the analysis.

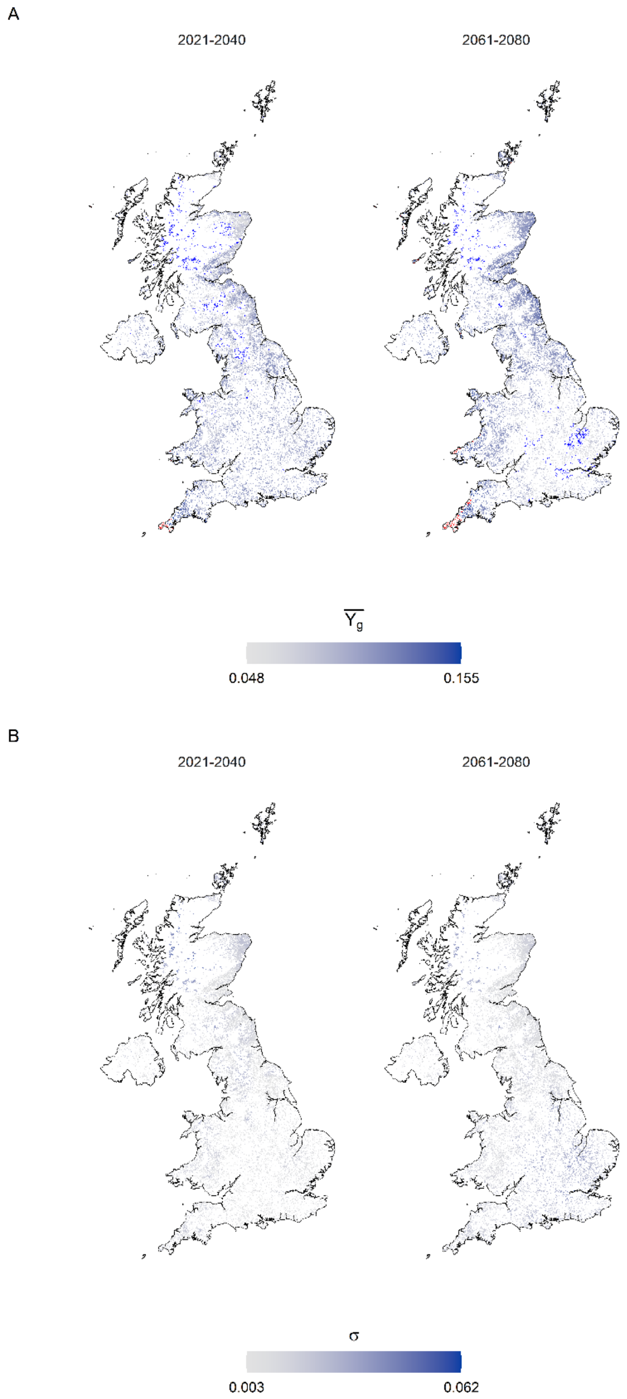


**Supplementary Fig. 8 |** Predicted future average yield gaps $Y_{gb}$ (**A**) in spring oats and their standard deviations σ (**B**) using 12 physics perturbations of climate model projections from UKCP Local Projections on a 5km grid. White grid cells contained no hosts and were excluded from the analysis. In panel **A**, values outside 1.5 times the interquartile range (IQR) above upper quartile and below lower quartile are shown in red and deep blue respectively.

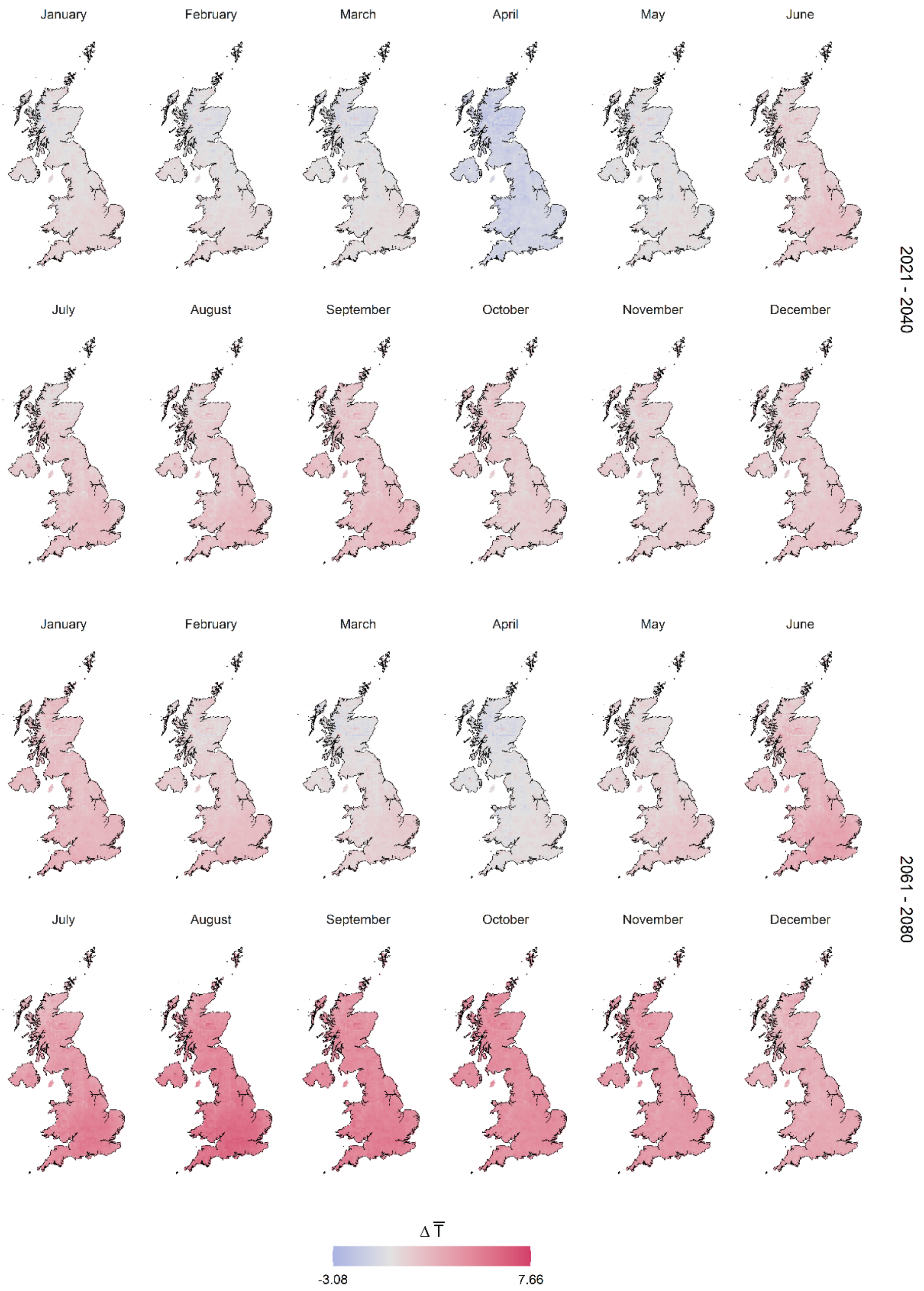


**Supplementary Fig. 9 |** Changes in future monthly temperature ΔT (°C) levels compared to current levels in the UK.

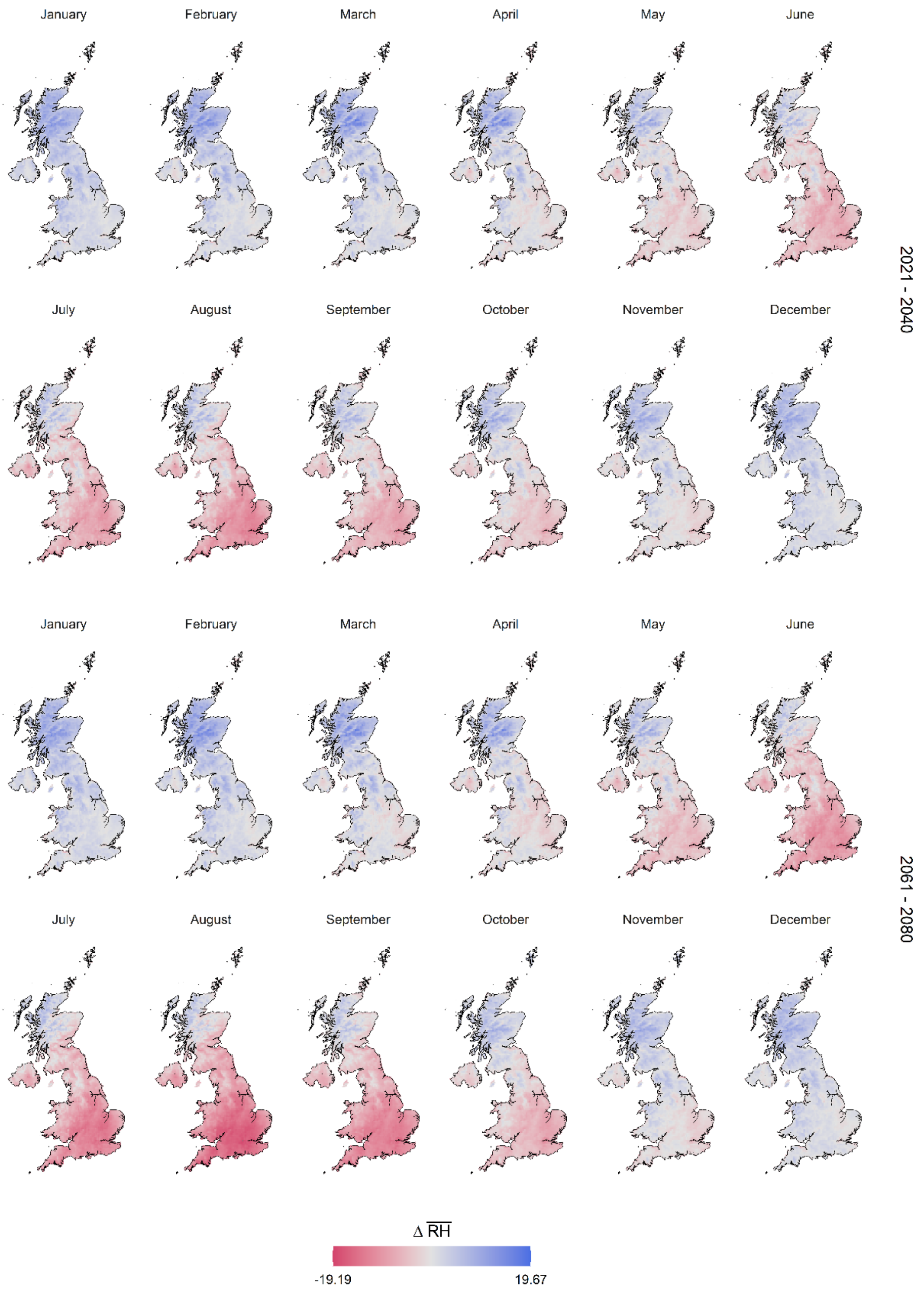


**Supplementary Fig. 10 |** Changes in future monthly relative humidity ΔRH (%) levels compared to current levels in the UK.

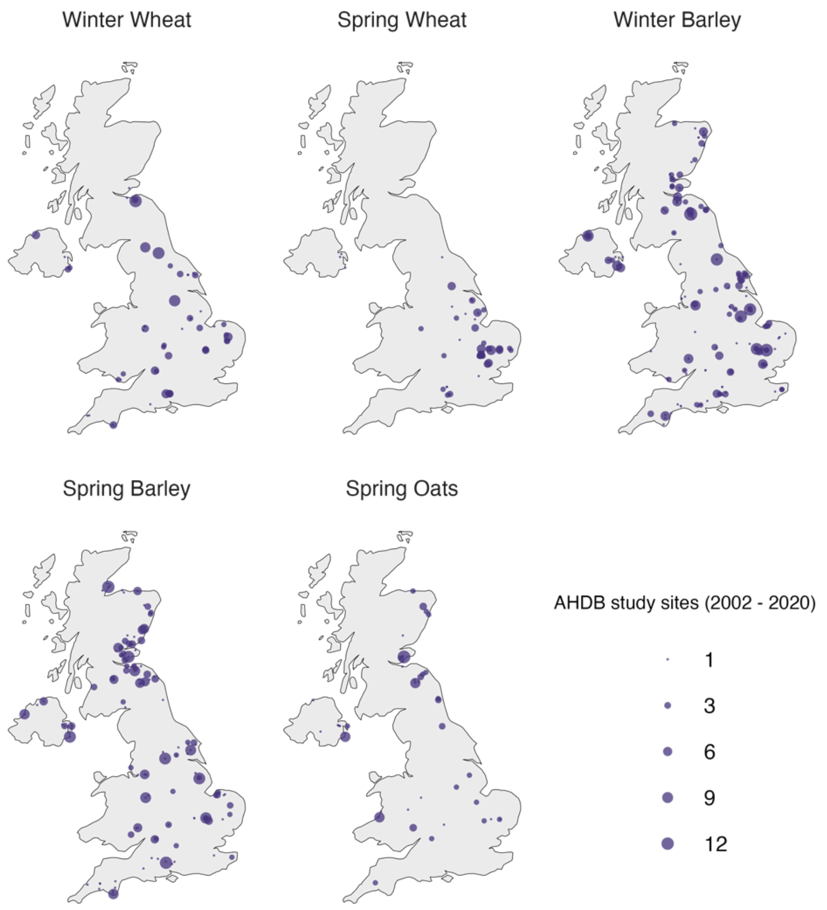


**Supplementary Fig. 11 |** Trial locations of AHDB crops studied in this paper. The size of point locations represent the number of years the experiments were conducted at those study sites.
