## Supplementary Tables for "Climate change, biotic yield gaps and disease pressure in cereal crops"

**Supplementary Table 1.** Count of AHDB studied crop varieties from 2002-2020. These are the varieties we used data for analysis.

| Year | Winter Wheat | Spring Wheat | Winter Barley | Spring Barley | Spring Oats |
|---|---|---|---|---|---|
| 2002 | 35 | 9 | 29 | 21 | 7 |
| 2003 | 45 | 10 | 0 | 0 | 0 |
| 2004 | 36 | 6 | 20 | 0 | 6 |
| 2005 | 38 | 7 | 23 | 26 | 5 |
| 2006 | 34 | 8 | 25 | 25 | 7 |
| 2007 | 36 | 6 | 22 | 21 | 11 |
| 2008 | 38 | 5 | 19 | 22 | 10 |
| 2009 | 34 | 6 | 17 | 21 | 11 |
| 2010 | 40 | 10 | 20 | 21 | 13 |
| 2011 | 40 | 9 | 20 | 22 | 17 |
| 2012 | 49 | 12 | 22 | 28 | 0 |
| 2013 | 54 | 12 | 26 | 33 | 14 |
| 2014 | 45 | 9 | 22 | 38 | 11 |
| 2015 | 51 | 13 | 21 | 24 | 12 |
| 2016 | 51 | 9 | 18 | 20 | 13 |
| 2017 | 46 | 9 | 20 | 21 | 12 |
| 2018 | 47 | 9 | 28 | 25 | 12 |
| 2019 | 39 | 10 | 29 | 26 | 12 |
| 2020 | 41 | 10 | 31 | 22 | 15 |
| **Total**† | **289** | **47** | **147** | **154** | **45** |

†Total unique varieties planted from 2002 – 2020 growing seasons.

**Supplementary Table 2.** Fitted parameter estimates of generalized-least-squares (GLS) to evaluate the effect of environment (time) and spatial locations (longitude and latitude) on treated yield ($Y_t$). Estimates in bold are statistically significant at 5% significance level. Values after ± area standard errors of estimates.

| | Constant | Year | Longitude | Latitude | ϕ§ | r† |
|---|---|---|---|---|---|---|
| Winter wheat | -105.22 ± 59.42 | 0.06 ± 0.03 | -0.07 ± 0.08 | 0.06 ± 0.09 | 0.22 | 0.25 |
| Spring wheat | 109.24 ± 69.95 | -0.05 ± 0.04 | -0.10 ± 0.16 | -0.07 ± 0.21 | 0.20 | 0.22 |
| Winter barley | **-168.98 ± 34.34** | **0.09 ± 0.02** | -0.06 ± 0.04 | 0.06 ± 0.04 | 0.23 | 0.35 |
| Spring barley | **-68.47 ± 32.04** | **0.04 ± 0.02** | **0.09 ± 0.04** | -0.01 ± 0.03 | 0.21 | 0.22 |
| Spring oats | -36.34 ± 65.30 | 0.01 ± 0.03 | -0.03 ± 0.09 | **0.31 ± 0.09** | 0.15 | 0.38 |

§Phi (ϕ) represents correlation between errors.
†Correlation (r) between observed and predicted $Y_t$ based on fitted GLS models.

**Supplementary Table 3.** Mean bootstrapped correlation coefficients of association between biotic yield gap ($Y_{gb}$) and weather variables from selected months. Values after ± are standard deviations of bootstrapped correlation coefficients.

| Crop | Temperature | Relative humidity |
|---|---|---|
| **Winter Wheat** | 0.42 ± 0.06 | 0.39 ± 0.06 |
| **Spring Wheat** | | 0.57$^{\dagger}$ ± 0.13 |
| **Winter Barley** | 0.32 ± 0.07 | 0.39 ± 0.06 |
| **Spring Barley** | 0.35 ± 0.08 | 0.35 ± 0.09 |
| **Spring Oats** | 0.30 ± 0.10 | 0.22 ± 0.14 |

$^{\dagger}$Correlation between $Y_{gb}$ and precipitation.

**Supplementary Table 4.** Fitted parameter estimates of generalized-least-squares (GLS) to study the effect of weather variables from selected months on biological yield gap ($Y_{gb}$). Estimates in bold are statistically significant at 5% significance level. Values after ± area standard errors of estimates.

| | Constant | Temperature | Relative humidity | $\phi^{\S}$ | $r^{\dagger}$ |
|---|---|---|---|---|---|
| Winter wheat | **-0.60 ± 0.17** | **-0.02 ± 0.003** | **0.01 ± 0.002** | -0.35 | 0.52 |
| Spring wheat | 0.04 ± 0.05 | | **0.002 ± 0.001**$^{¶}$ | 0.14 | 0.58 |
| Winter barley | **-0.64 ± 0.16** | **0.01 ± 0.004** | **0.01 ± 0.002** | 0.14 | 0.47 |
| Spring barley | **-0.52 ± 0.15** | **0.02 ± 0.005** | **0.006 ± 0.002** | 0.03 | 0.46 |
| Spring oats | 2.29 ± 1.32 | -0.39 ± 0.20 | -0.03 ± 0.02 | 0.41 | 0.41 |
| | | | **0.005 ± 0.003**$^{\ddagger}$ | | |

$^{¶}$Fitted coefficient of GLS between $Y_{gb}$ and precipitation.
$^{\ddagger}$Interaction of temperature and relative humidity.
$^{\S}$Phi ($\phi$) represents correlation between errors.
$^{\dagger}$Correlation (r) between observed and predicted $Y_{gb}$ based on fitted GLS models.

**Supplementary Table 5.** Current and future mean yield gap ($Y_{gb}$) and their 95% confidence intervals (CI) for the studied AHDB crops.

| Crop | 2002 - 2020 | | | 2021 - 2040 | | | 2061 - 2080 | | |
|---|---|---|---|---|---|---|---|---|---|
| | $\bar{Y}_{gb}$ | 95% CI | | $\bar{Y}_{gb}$ | 95% CI | | $\bar{Y}_{gb}$ | 95% CI | |
| | | LL | UL | | LL | UL | | LL | UL |
| **Winter Wheat** | 0.2142 | 0.2140 | 0.2143 | 0.2281 | 0.2280 | 0.2282 | 0.2425 | 0.2424 | 0.2426 |
| **Spring Wheat** | 0.2198 | 0.2195 | 0.2200 | 0.2038 | 0.2035 | 0.2041 | 0.1792 | 0.1790 | 0.1795 |
| **Winter Barley** | 0.1808 | 0.1806 | 0.1809 | 0.1867 | 0.1866 | 0.1869 | 0.1932 | 0.1931 | 0.1934 |
| **Spring Barley** | 0.1030 | 0.1029 | 0.1031 | 0.0746 | 0.0744 | 0.0747 | 0.0692 | 0.0690 | 0.0693 |
| **Spring Oats** | 0.1198 | 0.1195 | 0.1200 | 0.0972 | 0.0968 | 0.0976 | 0.1013 | 0.1010 | 0.1017 |

**Supplementary Table 6.** Median and interquartile ranges (IQR) of standard deviations of the predicted future average yield gaps ($Y_{gb}$) using 12 physics perturbations of climate model projections from UKCP Local Projections on 5km grid.

| Crop | 2021 – 2040 | | 2061 - 2080 | |
|---|---|---|---|---|
| | Median | IQR | Median | IQR |
| **Winter wheat** | 0.0129 | 0.0029 | 0.0152 | 0.0024 |
| **Spring wheat** | 0.0129 | 0.0029 | 0.0152 | 0.0024 |
| **Winter barley** | 0.0107 | 0.0032 | 0.0142 | 0.0027 |
| **Spring barley** | 0.0132 | 0.0040 | 0.0137 | 0.0038 |
| **Spring oats** | 0.0108 | 0.0082 | 0.0150 | 0.0112 |
